# Signals of microbial growth learned from single amplicon samples

**DOI:** 10.1101/2023.11.07.565791

**Authors:** Philippe Chlenski, Deirdre Ricaurte, Itsik Pe’er

## Abstract

Irregularities in metagenomic whole-genome shotgun (WGS) read coverage can arise in quickly replicating microbial populations. These irregularities, summarized as peak-to-trough ratios (PTRs), are correlated with growth rates. This study seeks to explore the presence of similar coverage irregularities in 16S amplicon datasets, where multicopy diverged 16S genes provide an opportunity to explore coverage at different positions on the bacterial chromosome. To this end, we propose a model of Operational Taxonomic Unit (OTU) observations under replication and sequence similarity, from which we derive a method for simultaneous copy number correction and dynamics estimation by gradient descent. We conduct a series of benchmarks on synthetic data, determining a set of heuristics for when such methods may be applied, and compare our method with WGS-based methods on a real dataset. We find no correlation between coPTR estimates and our method, suggesting further modifications to our method may be required.

## 1 Introduction

### 1.1 Peculiarities of 16S amplicon sequencing

Amplicon sequencing of the 16S ribosomal RNA gene is one of the key techniques in analyzing microbiomes. It is popular in large part because the diversity of the 16S gene is reasonably well understood, and analysis is streamlined by the short length of the regions being amplified (avoiding the need for contig assembly) and the existence of reference databases such as SILVA[17] and Greengenes[4]. These 16S sequences are commonly clustered into operational taxonomic units (OTUs), a coarse-grained bin corresponding to all sequences within some level of nucleotide identity (commonly 97%) of one another.

On the other hand, 16S amplicon sequencing suffers from several shortcomings compared to whole-genome shotgun (WGS) methods: at the inter-genome level, 16S gene sequences can fail to distinguish organisms at many taxonomic levels [12]. At the intra-genome level, the 16S gene can be both multicopy and diverged [2, 19]. Multicopy 16S genes can produce OTU counts that are several times higher than what would be consistent with the actual species abundances, creating a need for copy number correction [6, 1, 10]. Diverged 16S sequences, on the other hand, can look like correlated but distinct OTUs. Several studies have looked explicitly at 16S copy number variation and its connection to growth rate by way of ecological adaptation [16, 21, 18, 7].

Finally, amplicon datasets provide little insight into the chromosomal content of the species in the sample. They may be unable to distinguish between structural variants for genes distinct from the markers being surveyed, and amplicon reads can only be mapped to very small subregions of the entire reference genome.

### 1.2 Peak-to-trough ratios

In most Prokaryote species, replication proceeds bidirectionally from a single origin of replication (OOR) and ends at the replication terminus (RT) at the opposite end of the chromose. WGS reads from actively replicating species exhibit a log-linearly decreasing coverage pattern from the OOR to the RT. This pattern can be thought of as the limit of the sum over copy numbers for many chromosomes in partial states of replication. The coverage ratio between the OOR and the RT, called the peak-to-trough ratio (PTR), can be shown to be correlated to bacterial growth rate [8].

Since classical PTR estimation looks at irregularities in WGS read coverage across the entire chromosome to estimate the rate of replication in bacterial species, this analysis cannot straightforwardly be applied to marker genes. However, the multicopy diverged nature of the 16S gene offers us an opportunity to seek out signals of replication in amplicon sequences: if two 16S genes can be distinguished by their sequence, and if their positions are known relative to that genome’s OOR, then in principle this is sufficient to fit an entire coverage curve consistent with microbial growth rate estimation.

### 1.3 Contributions

In this paper, we propose a mathematical framework for inferring PTRs from 16S amplicon data. By explicitly modeling the expected coverage at each position in the genome and binning genes according to their sequence similarity, our method simultaneously corrects for 16S copy numbers and estimate microbial growth rates.

Additionally, we conduct a number of benchmarks on synthetic and real data. We derive coverage bounds within which our method can learn PTRs with reasonable accuracy and experiment with various aspects of our model. We compare to coPTR, the leading method for PTR inference in WGS data, on simulated and real data.

We find that our method works well in simulated cases, but fail to demonstrate effectiveness in existing datasets. We stipulate that this is largely a function of dataset quality and outline experimental validation techniques that can determine the empirical usefulness of our technique more explicitly.

## 2 Methods

### 2.1 Reference data collection

To generate the database of viable specimens, we downloaded the sequences and genomic locations of the 16S rRNA and dnaA genes for all PATRIC [13, 3] representative genomes. Then we filtered them as follows:

1. Truncate reference 16S sequences according to the primer used in the study (e.g., keep only the V3-V4 hypervariable regions of the gene)
2. Remove genomes without any contig that has multiple distinct 16S sequences
3. Remove genes that share a 16S sequence with any of the genomes removed in step (1)
4. Remove any additional genomes that fail criterion (1) after step (2)
5. Repeat steps 1–3 until convergence
6. Keep only genomes with a single known OOR

This series of strict filters guarantees that each 16S sequence that appears in the database is only known to appear on contigs with multicopy diverged 16S genes and is not thought to be confounded by any singleton RNA signals.

It can also be helpful to incorporate priors as a preprocessing step. For instance, when working with a sample that is known to contain a limited set of species, pre-filtering the database can result in a richer set of genomes for testing as fewer are removed to avoid confounding in step (2).

### 2.2 Generative model

All generation and inference code is implemented in PyTorch [14].

#### 2.2.1 Notation

Let *S, T, G*, and *N* be the number of samples, OTUs, genes, and distinct 16S rRNA gene nucleotide sequences in an experiment. Let *s, t, g*, and *n* be indices for specific samples, OTUs, genes, and sequences. So, for instance, an OTU table would be represented by a *T*×*S* matrix **Y**, and *Y*_*t,s*_ would be the count of reads mapping to OTU *t* in sample *s*.

#### 2.2.2 Modeling coverage

Under exponential growth assumptions, the log ratio of reads from the OOR to reads from the RT is summarized in the matrix **R**, where

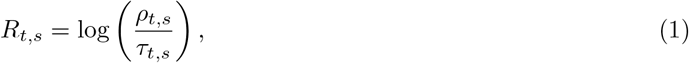

and *ρ*_*t,s*_ is the replication time and *τ*_*t,s*_ is the generation time. Coverage decays log-linearly from the OOR to the RT at a rate set by the PTR.

For a known PTR, let *i* be the total number of bases in a microbial genome and *j* be the locus for the OOR. We compute *δ*_*t*_(*l*), the distance of locus *l* from the OOR *l*_0,*t*_ on a circular chromosome of size *L*_*t*_, as

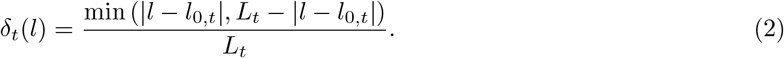

Then, letting **A** be the *T*×*S* matrix of abundances (defined as coverage at the OOR), we model coverage at locus *l* as

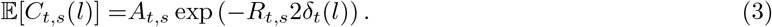

These expectations can be used directly as the *λ* parameter of a Poisson distribution to model read count expectations. Alternatively, they can be transformed into read probabilities by a simple normalization:

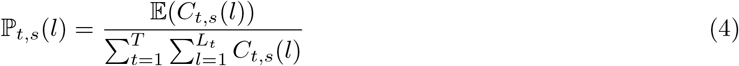

#### 2.2.3 Determining OTU read probabilities

Let **A** and **R** be *T* × *S* matrices of microbial relative abundances and log-PTRs, respectively. Letting *l*_*g,t*_ denote the locus at which gene *g* starts in the genome of OTU *t*, we additionally define an *N*×*G* sequence-sharing matrix **S**, a *G*×*T* distance-to-OOR matrix **D**, and a *G*×*T* gene-genome membership matrix **M**.

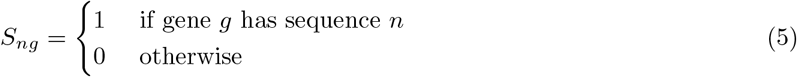

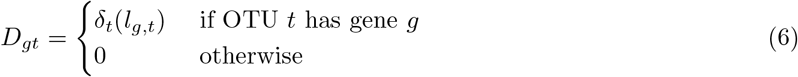

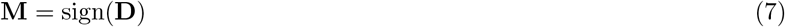

The matrices **S, D**, and **M** fully describe the way a sample with known abundances and growth rates will behave. Putting this all together, we can produce the *N* × *S* coverage matrix **Ŷ**:

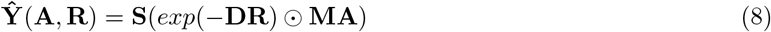

#### 2.2.4 Likelihood model for read counts

It is natural to adopt a Poisson likelihood function for unnormalized count data. Since the entries of matrix **Ŷ** already correspond to expected read counts, they can be used directly as a Poisson likelihood function parameter. Let **Y** refer to the *N* × *S* empirical OTU count matrix.

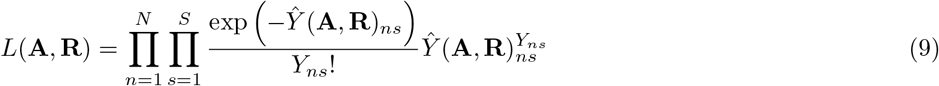

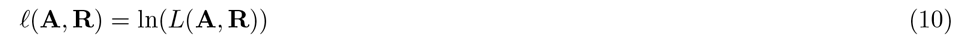

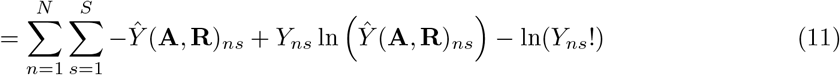

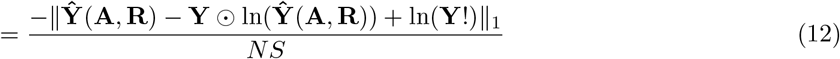

Negating Eq. 12 yields the negative log-likelihood, which is used in practice for optimization.

### 2.3 Inference model

#### 2.3.1 General formulation

A matrix of empirical observations **Y** and matrices **S, D**, and **M** reflecting our knowledge of the distribution of 16S RNA sequences, plus some loss function 𝔼(**Ŷ**, **Y**), can also be used to estimate unknown abundance and log-PTR matrices **A** and **R**. In other words, we are trying to find approximations **Â**, 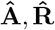 such that we attain:

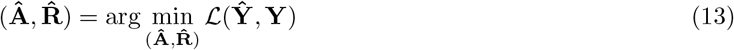

This problem can be solved using gradient descent. The general form of the gradients is:

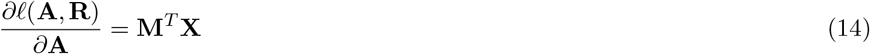

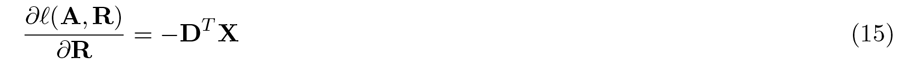

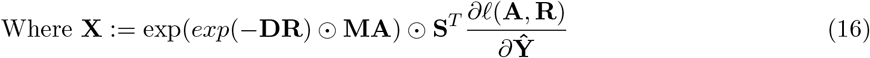

#### 2.3.2 Mean squared error loss

When ℒ(**Ŷ**, **Y**) is a mean squared error loss function (corresponding to a gaussian likelihood model), then the gradients are

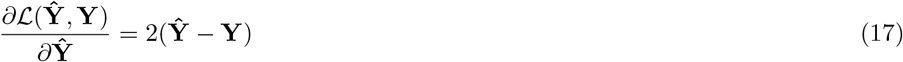

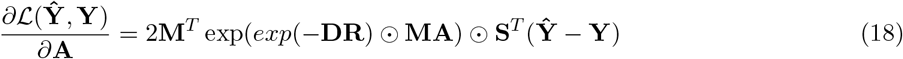

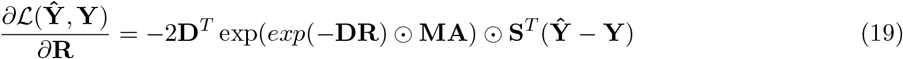

#### 2.3.3 Poisson negative log-likelihood loss

Let ⊘ denote elementwise division. When ℒ (**Ŷ**, **Y**) is the negative Poisson log-likelihood (the negation of Eq 13) the gradients are

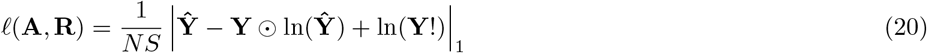

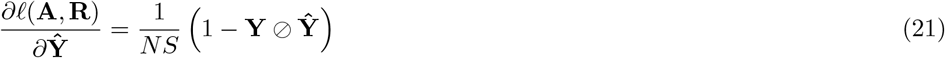

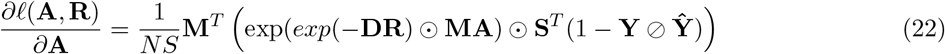

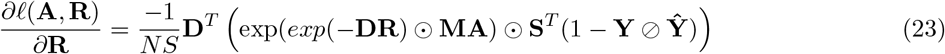

### 2.4 Further optimizations

#### 2.4.1 Regularization

Regularization is straightorward to apply to this model, as an additive term to the loss function. For instance, applying an *L*_1_ penalty to the values of the estimated abundance matrix **Â** encourages sparsity, which can be consistent with patterns of complete species absence across samples in microbiome datasets.

#### 2.4.2 Clipping and filtering estimates

Another way of incorporating domain knowledge into the model is to clip the values of the log-PTR estimates 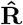, for instance, to the interval [0,1]. This has the desirable effect of keeping all PTRs within biologically plausible bounds, which can force the system out of consistent but nonsensical configurations of log-PTRs, and in particular avoids the problem of inverted coverage curves corresponding to negative log-PTRs.

#### 2.4.3 Modeling amplicon bias

Amplicon bias is a phenomenon that can seriously distort OTU abundances, but it is not well-characterized in a way that lends itself to explicit, reference-based correction. [11] Instead, we can introduce an amplicon bias term directly into our model. We assign each OTU its learnable bias parameter but assume that biases are consistent across samples. Thus, we have an *N* -length bias vector **h**, and we model

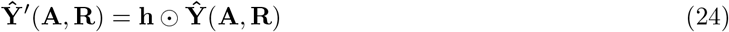

where ⊙ means each column vector of **Ŷ** is multiplied by **h**.

**h** is learnable by gradient descent, but this approach is only theoretically justified when *S* is larger than *N*, so that the system is not underdetermined. In practice, the empirical investigation into the effects of modeling amplicon bias in Section 3.4 suggests that this requirement can be relaxed, however.

### 2.5 Data generation and processing

#### 2.5.1 Synthetic data

The likelihood model in Eq. 3 specified how a system of *T* OTUs across *S* samples with known abundances, log-PTRs, and distributions of 16S sequences can be used to derive expected read counts for *T* OTUs in *S* samples.

To model OTU counts, we took real genomes from the database constructed in Section 2.1, restricted to the subset of complete (single contig) genomes, and simulated reads by drawing from a Poisson distribution parameterized by the expected read-count matrix **Ŷ** (**A, R**) for known but held-out **A** and **R**. We used the resulting (sampled) OTU table directly as amplicon sequence data.

To model a greater diversity of possible microbial systems, we introduce an optional sparsity term *b*, such that a *T*×*S* sparsity matrix **B** is drawn with each entry corresponding to an independent Bernoulli trial with probability *b* of success. Multiplying **A** by **B** encourages sparsity in the simulated system.

For finer-grained control over the read expectations, we also introduced a convenience multiplier term *m*, referred to as the “scale” variable. This can be thought of as the total amount of time spent on sequencing: thus, it is semantically different from multiplying all absolute abundances by a scalar factor, although the actual effect on the expectations is the same.

To generate whole-genome shotgun (WGS) data, we used the single-sample variation of the simulation framework to sample each possible position in the genome. To generate reads, we first downloaded the corresponding FASTA files from PATRIC. Then, for each observation count at each locus, we drew the corresponding number of 300-base reads from the FASTA file. The full list of reads was then shuffled and concatenated into one FASTQ file. This single-sample process was repeated *S* times to generate a full set of samples.

To control the ratio of WGS to amplicon reads, we introduce an optional downsampling term *p* such that WGS reads are only kept with probability *p*, whereas all amplicon counts are kept with probability 1 so long as they fall within a 16S RNA region.

#### 2.5.2 Pipeline for processing 16S data

We used VSEARCH[15] to process FASTQ reads from amplicon sequences into deduplicated FASTA files with per-sample multiplicities. After trimming the reference sequences according to the primers used in the study, we used VSEARCH with an exact identity requirement (--id 1.0) to enforce exact matches to reference sequences.

A second phase of OTU table postprocessing checks the OTU sequences against the reference database. OTUs are kept only if candidate genomes are found such that each OTU corresponding to that genome has nonzero multiplicity in the OTU table. Finally, the filtered OTU table is run through a script that performs the optimizations laid out in Section 2.3 and outputs tables corresponding to the estimated abundances **Â**, log-PTRs 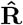, estimates of the number of reads used per cell, and amplicon bias terms **h** if applicable.

#### 2.5.3 Processing WGS data

To infer PTRs on metagenomic data, we use coPTR[5] with several modifications. First, for the read mapping portion of the coPTR pipeline, we created a special database of whole genome sequences for the reference genomes in our database. This ensures that IDs match during comparison and speeds up processing by avoiding mapping to genomes that do not have multiple diverged 16S RNA sequences.

For comparing coPTR on real data, we used the DIABIMMUNE dataset [22, 9, 20], restricted to the 1,009 samples for which WGS and 16S amplicon sequencing were performed in parallel. The corresponding NCBI SRA accessions are PRJNA290380, PRJNA231909, and PRJNA290381.

## 3 Results

### 3.1 Poisson vs. gaussian likelihood

We analyzed errors in PTR estimates for synthetic datasets with 10 genomes, 20 samples, and varying scaling factors *m* using both Poisson (blue) and Gaussian likelihood functions (mean squared error; red) to score estimates. A boxplot of experiment results is shown in the right half of Fig. 1.

**Figure 1:**
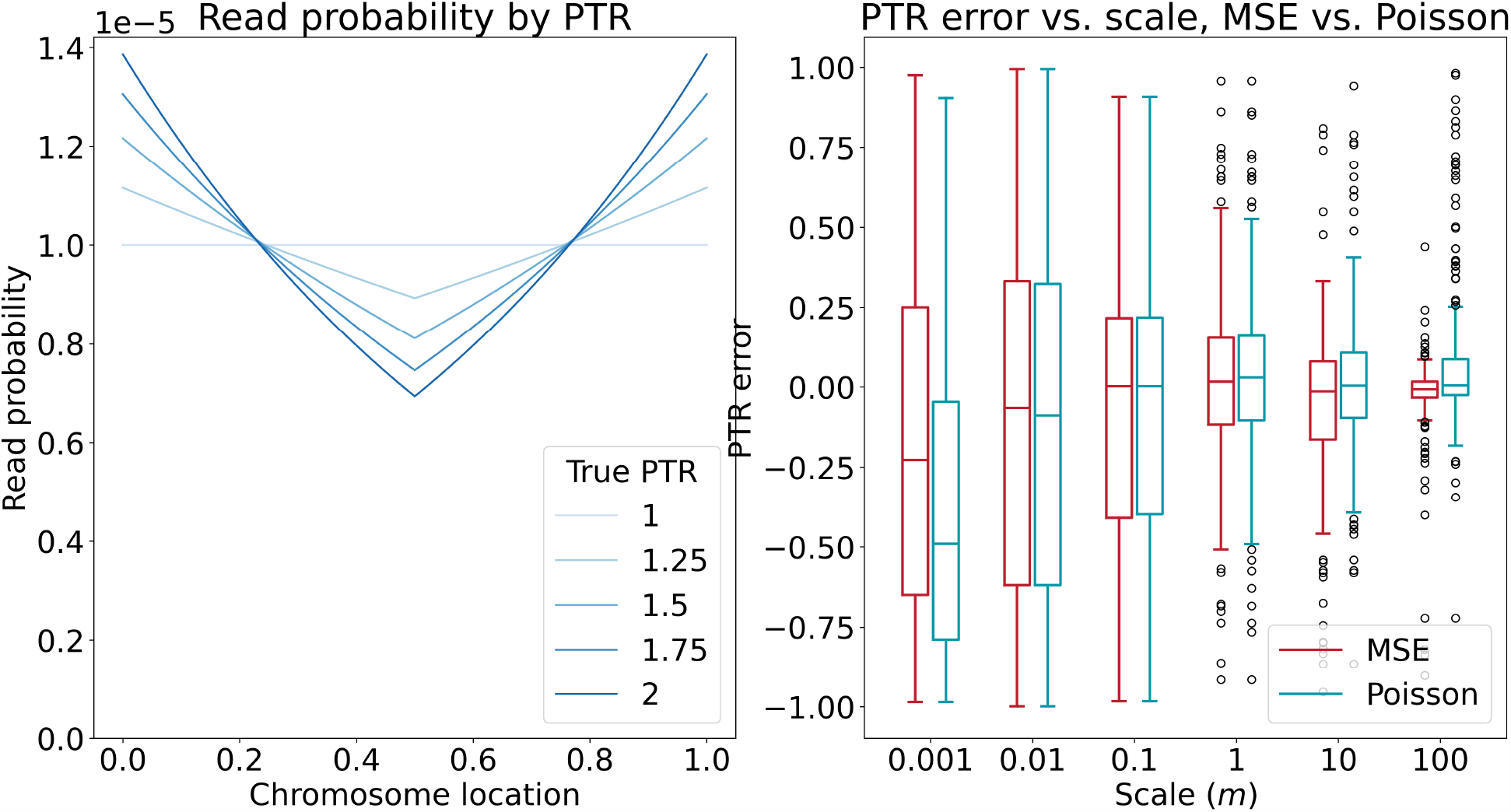
Left: Read probabilities decay log-linearly from the OOR (at x = 0) to the RT (at x = 0.5). Perposition read probabilities are shown for a 100,000-base chromosome for different PTRs. Right: There is no substantial difference between Poisson and MSE-based regression approaches, although Poisson regression is more well-motivated in the biological context and matches the modeling approach used to generate synthetic data. There may be some performance decrease for very large scaling factors, but these correspond to coverage depths much larger than those typically encountered in amplicon datasets.

We found no appreciable difference in errors in most parts of the range, except for when the scaling factor was 100. In this case, the Gaussian likelihood outperformed the Poisson likelihood model. This is an interesting observation, but may be of limited practical use since (according to the number of reads plotted in the right half of Fig. 2) this would yield more than 10,000 reads per OTU per sample—a sequencing depth rarely seen in actual experiments.

**Figure 2:**
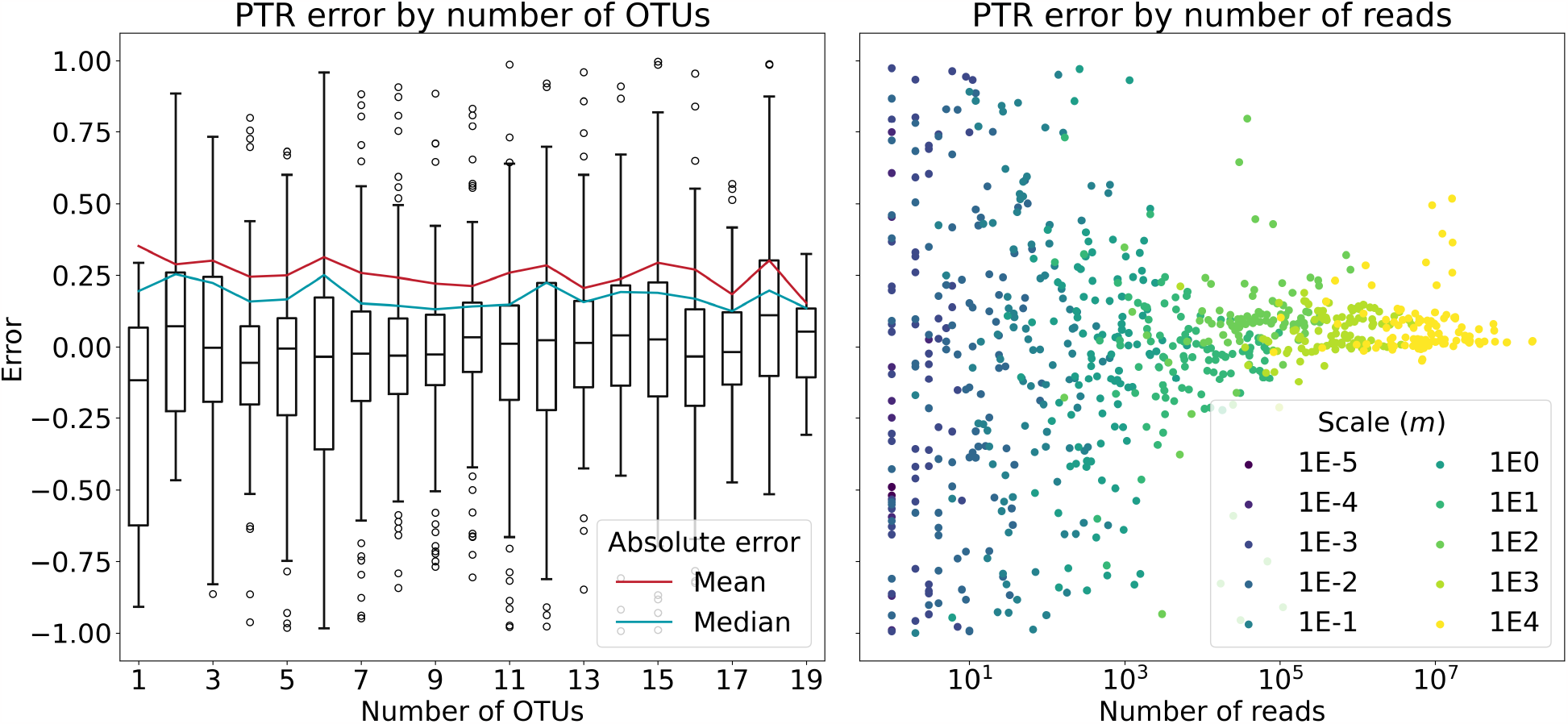
**Left:** No trend in PTR estimation error is observed as the number of OTUs increases. **Right:** Error goes down substantially with number of reads. This suggests that estimates with at least 1,000 hits per OTU per sample can be analyzed with high precision.

### 3.2 Effect of number of OTUs

We sampled a synthetic dataset with abundances of 100 per OTU and log-PTRs drawn from a uniform distribution on the interval [0, ln(2)], varying the total number of OTUs simulated and tracking the average error in each log-PTR estimate in 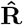. The effect of the number of OTUs on error is shown in the left left of Fig. 2.

There was no appreciable trend in mean (red) or median (blue) error as the number of OTUs increased, nor was there any trend in the overall distribution of errors. In total, it appears that simultaneously estimating PTRs for a large number of OTUs, as one might encounter in real microbial ecosystems, causes any decline in predictive accuracy.

### 3.3 Effect of number of reads

We analyzed the errors in PTR estimates for a synthetic dataset with 5 OTUs, 20 samples, and uniformly drawn log-PTRs as we varied the expected read count by changing the multiplier term *m* for the expected read counts. The results of this experiment are shown in the right half of Fig. 2.

We found that, for very small multipliers, predictive performance was no better than random. However, performance improved greatly with higher read count expectations. We found that total substantially constricted around 1,000 reads per OTU per sample, suggesting this is a reasonable minimum number of hits to perform PTR estimation. Past this minimum, improvement in predictive performance was negligible.

### 3.4 Amplicon bias modeling

We explored estimate performance for Poisson regression on systems of 20 samples and 10 OTUs over various expected value multipliers. We modeled amplicon bias according to the method described in Section 2.4.3, by applying an OTU-specific bias matrix identically to each sample and each column of the estimated abundances **Ŷ** (**A, R**). We used a solver with a free *N* -length bias term both with and without actual bias in the system. The results for this experiment are summarized in Fig. 3.

**Figure 3:**
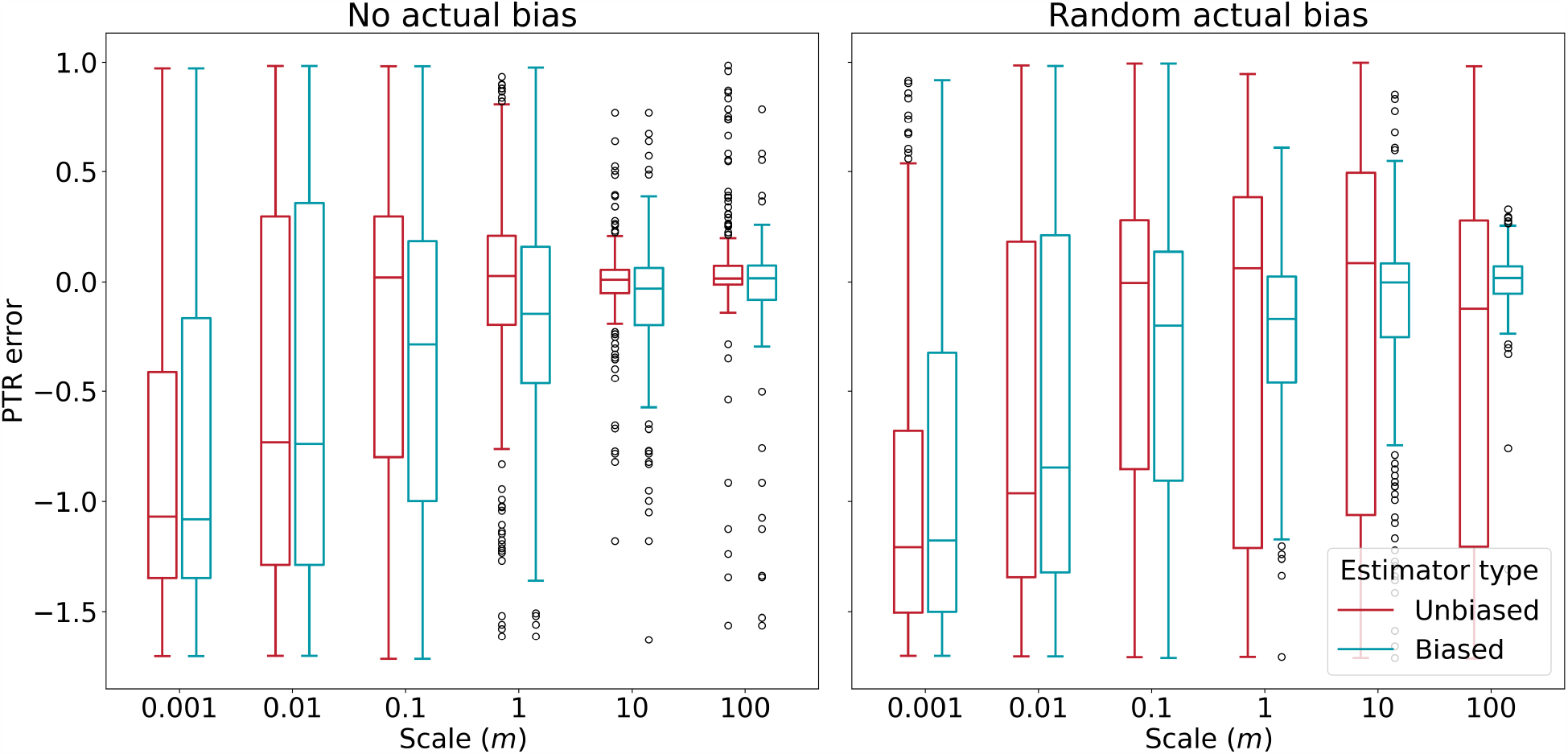
**Left:** Without any actual amplicon bias in the modeling process, an estimator with a free bias term does not do substantially worse than an estimator without a free bias term. **Right:** On the other hand, when amplicon bias is built into the modeling process, an estimator with a free bias substantially outperforms an estimator without a free bias term.

In both figures, red boxplots represent an unbiased estimator, i.e. one which does not have a learnable amplicon bias term. Blue boxplots represent identical estimators, except that they do have learnable amplicon bias terms. In the left half of Fig. 3, both estimators are used in the absence of actual simulated amplicon bias with no detectable difference in performance. This indicates that correcting for amplicon bias when it is not needed does not degrade model performance.

On the other hand, the left half of Fig. 3 shows the performance of these same classifiers in a system where amplicon bias has been simulated. Here, the biased estimator achieves substantially lower error than the unbiased estimator. Thus, although there appears to be relatively little risk in accounting for bias when it is not needed, it is catastrophic not to account for bias when it actually is present in the system.

### 3.5 Comparison with coPTR

#### 3.5.1 Synthetic data generation

In order to evaluate the performance of our proposed method against coPTR, synthetic datasets were generated. The synthetic read counts were drawn from Poisson distributions, with parameters determined by the genome size, abundance, and PTR as elaborated in Section 2.5.1. Three distinct systems were sampled, characterized by multipliers *m* ∈ {0.1, 1, 10} and a sparsity matrix **B** with an expected density *b*. The synthetic data generation was conducted with the following parameters:

- Number of genomes per sample: 20
- Number of samples: 10
- Sparsity matrix elements *B*_*t,s*_∼*Bernoulli*(*b*)
- Abundance matrix elements *A*_*t,s*_∼*Exp*(1)
- log-PTR values *R*_*t,s*_∼*Uniform*(0, ln(2))
- Read counts *Y*_*t,s*_∼*Poisson*(*m***B**⊙𝔼[*C*_*t,s*_(*l*)]), as in Eq. 3.
- WGS reads were downsampled uniformly by a factor of 100

We found that our method beats coPTR at lower read counts, but even with downsampling, the amount of reads coPTR had available tended to be much higher. These results are shown in Fig 4.

**Figure 4:**
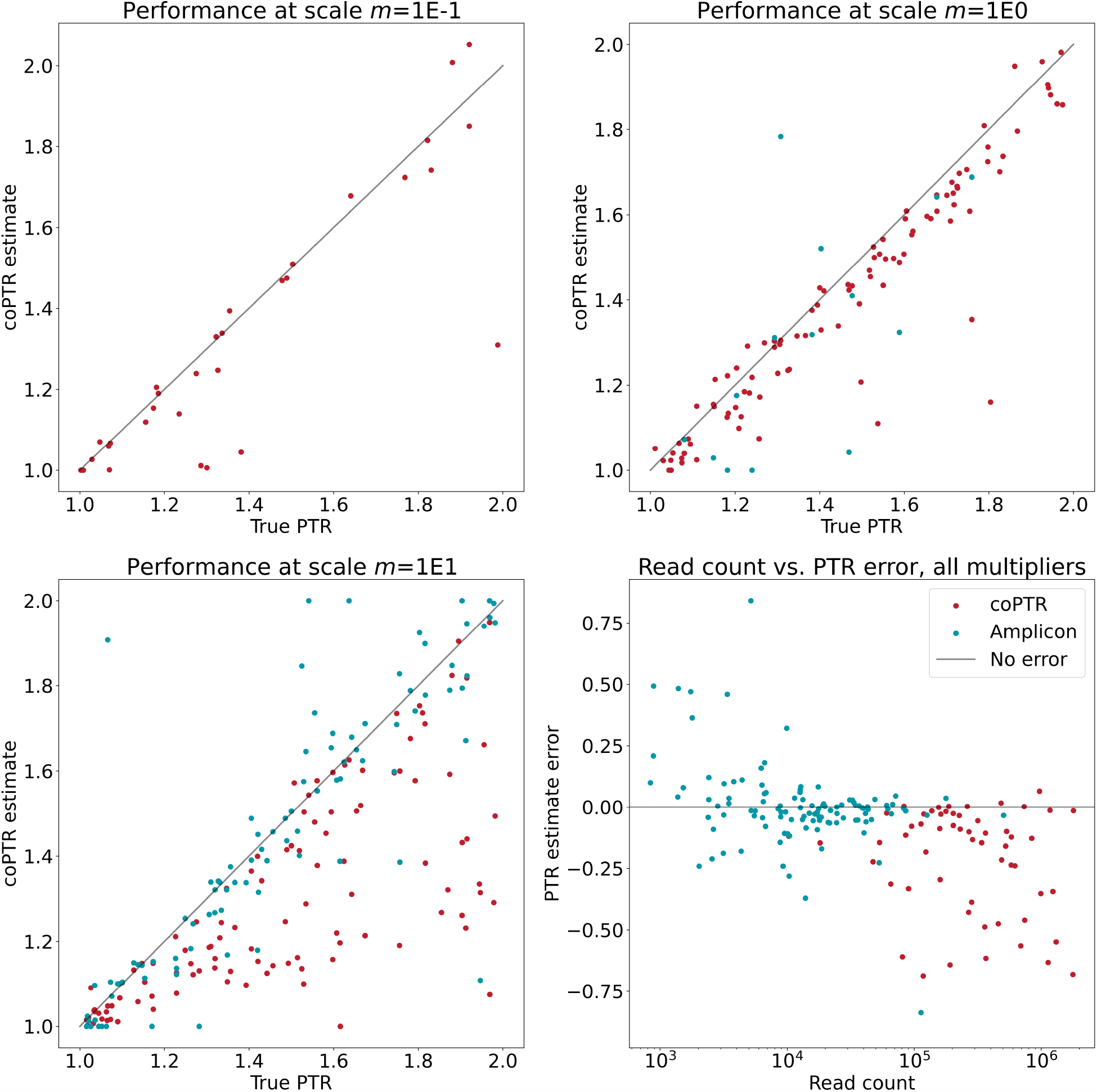
**Top left, top right, bottom left:** A comparison of coPTR to our method for synthetic data. We see that even when downsampled so that there is 100-fold more coverage on 16S regions than other areas of the chromosome, coPTR outperforms our method at low count expectations. This is most likely driven by 16S regions covering less than 1% of the chromosome, causing there to be far more WGS reads overall. **Bottom right:** We aggregate the other three figures and plot the relationship between read count and error. This shows that, when controlled for the overall number of reads per genome, our method actually outperforms coPTR on synthetic data.

In Fig 4, we show scatterplots of true versus predicted PTRs for both coPTR and our method at scaling factors of *m* = {0.1, 1, 10}. For *m* = 0.1 (top left), we did not produce any estimates. For *m* = 1 (top right), we produce some low-reliability estimates. However, at *m* = 10 (bottom left), we noticeably outperform coPTR.

In the bottom right plot of Fig 4, we summarize the total number of reads per OTU per sample used to generate each estimate, pooled across all *m* values. This figure reveals that coPTR always uses more reads than our method, even when comparing coPTR with *m* = 0.1 and our method with *m* = 10. This partially explains the performance gaps at lower expectations, but also reveals that we can outperform coPTR when controlling for the number of reads.

#### 3.5.2 DIABIMMUNE dataset comparison

Finally, to determine the accuracy of our method on real data, we explored the correlation between coPTR estimates and our predictions on the DIABIMMUNE dataset. The 16S component of the dataset consisted of paired-end reads from the V4 hypervariable region sequenced on the Illumina HiSeq 2500 platform.

First, we ran the 1,009 samples, which had both WGS and 16S reads available through a VSEARCH pipeline for *de novo* OTU table creation. Then, we applied our inference pipeline with the amplicon bias term and without PTR clipping or regularization. Finally, for the 78 samples for which we produced any estimates satisfying our confidence criteria, we produced coPTR-based PTR estimates using the method described in Section 2.5.3.

We found that there was no correlation between our method and the estimates produced by coPTR (Spearman *R* = 0.01, Pearson *R* = 0.08). A scatterplot of our estimates compared to coPTR’s is found in Fig 5. This figure not only shows our lack of correlation to coPTR, it also reveals a tendency to wildly overestimate PTRs in the absence of clipping, with the top estimated PTR being over 16 million. Other combinations of hyperparameters (e.g. turning off amplicon bias modeling, increasing regularization, etc) produced similarly uncorrelated results.

**Figure 5:**
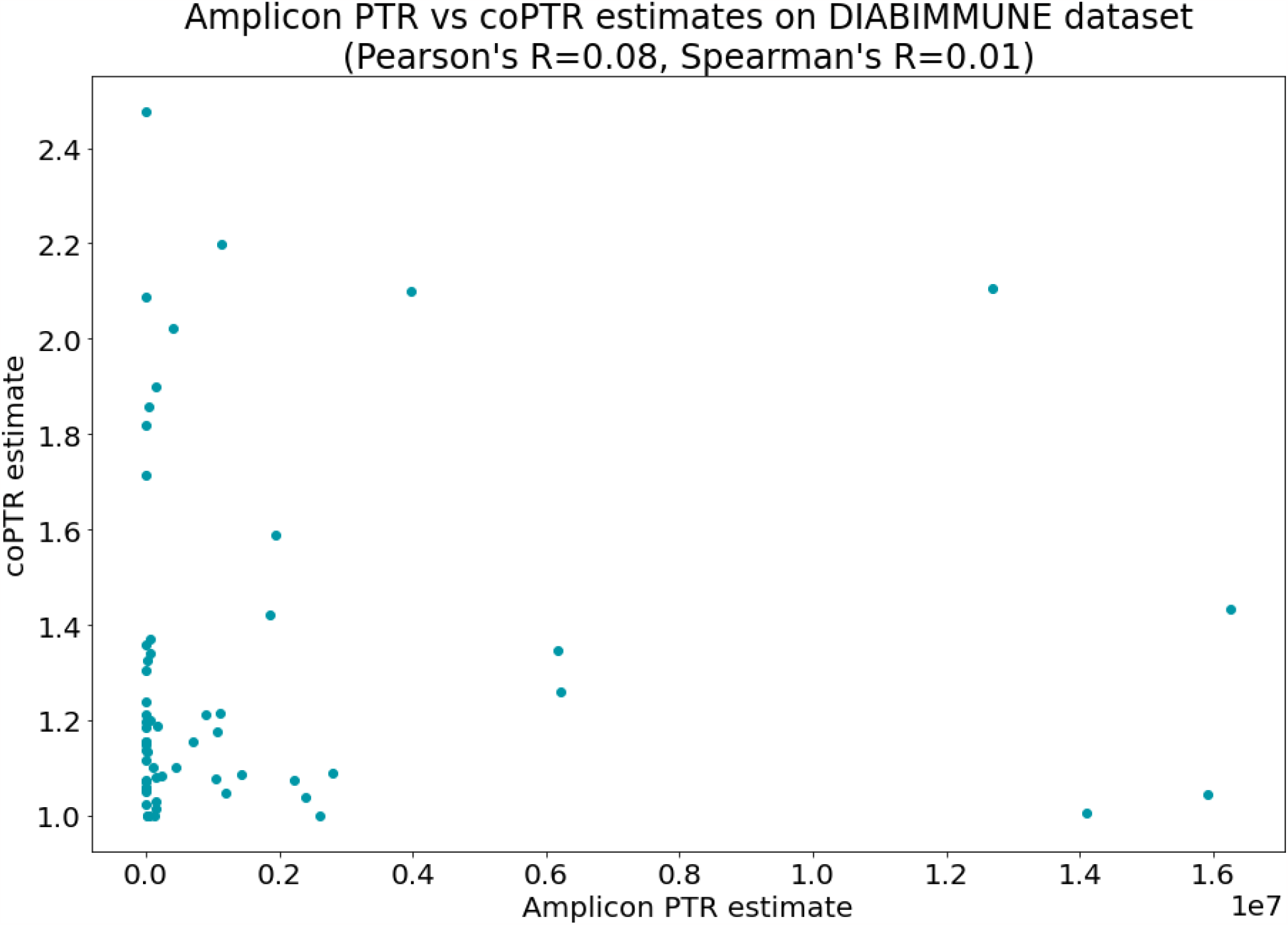
No correlation between coPTR PTR estimates and our amplicon estimates. In particular, our method tends to greatly overestimate PTRs, sometimes by many orders of magnitude. This may be driven by a mis-specification of how actual noise looks, low read coverage, or other factors.

## 4 Discussion

While our method effectively recovered PTRs under synthetic conditions matching our generative hypothesis, we could not find a dataset to observe the corresponding coverage patterns. We present an incomplete list of reasons why this may have been the case:

1. **Inadequate coverage**. For most genomes in most samples, there was insufficient coverage to produce accurate estimates of PTRs. Our investigation into synthetic data suggests that several thousand reads should be sufficient, but other sources of non-sampling-based noise may increase coverage demands in empirical contexts. In the future, deeper sequencing of communities may bring read counts within an acceptable range.
2. **Short reads**. Most 16S datasets are sequenced using amplicon primers that amplify short subsequences of the 16S gene such as the V4 hypervariable region used in the DIABIMMUNE study. This
3. substantially constricts our method’s resolution by reducing the effective database size and increasing the probability of contamination from identical sequences in non-target genomes. While our method is theoretically robust to gene sequences observed in multiple genomes, it makes the nontrivial assumption that we have a complete catalog of genomes to which a given read can map. As longer-read sequencing technologies become more common, more studies sequencing the entire V1-V9 region of the 16S gene may appear. These would be especially useful for testing our model.
4. **Dataset complexity**. Gut communities are quite complex, with several orders of magnitude difference in abundances and extensive strain-level variation. This could create variation patterns that disrupt the signal we are looking for. Simple synthetic datasets, such as amplicon sequences of simple *in vitro* communities, might be better for a proof of concept.
5. **Confounding by evolutionary strategy**. As mentioned earlier, 16S copy number has been associated with different evolutionary strategies at the strain level. It is reasonable to assume that evolutionary strategies may also have implications for growth rates, and therefore true PTRs, in empirical case studies, and that this is inadequately controlled for in our study. In the absence of an explicit model of the relationship between variables, it is not clear how these variables can be controlled for in a principled way.
6. **Amplicon bias**. We use a naive model of amplicon bias based on *in silico* performance. This may fail to model the more complex ways in which amplicon bias actually manifests.
7. **Poor reference data**. Our reference data is approximate, and some of it is derived from metagenome assemblies. It is quite likely that there are some errors in our references or that we have inadequate resolution to capture the full range of 16S copy number variation at the OTU-bin level.
8. **Copy number variation below the strain level**. If 16S sequences are unreliable predictors of copy number, our ability to model 16S coverage collapses.
9. **Correlation with coPTR**. Since coPTR estimates are also approximate, we lose some insight into true microbial dynamics by comparing our estimates with coPTR.

It is also possible that the coverage model we are using is flawed, imprecise, or based on wrong assumptions. We have been unable to demonstrate the anticipated patterns of growth dynamics in the data available to us. Still, we present this method to enable more conclusive experimental validation in the future.

## Supporting information

MLCB 2023 Extended Abstract

## Appendices

### A Lagrange model of bin coverage

The first approach we developed to solving this problem cast it as a constrained optimization problem. For a single genome, we wanted to find

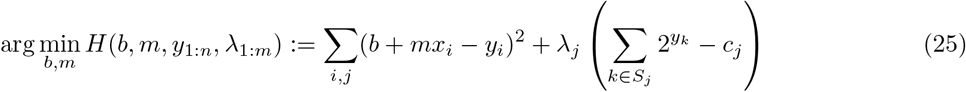

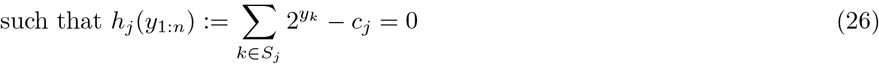

Where *x*_*i*_ is the genomic coordinate of some 16S gene, *y*_*i*_ was its coverage, and the Lagrange constraint forced coverages for all genes sharing a sequence (represented as the set *S*_*j*_) to sum to *c*_*j*_, the observed OTU coverage. This formulation allows us to determine *m* and *b*, the slope and intercept of the log-coverage, from which the PTR could be computed as

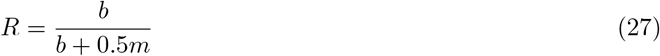

For multiple genomes, we can expand this to:

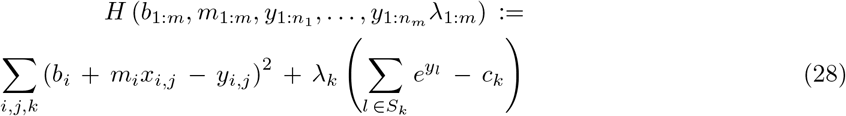

Subject to the same constraints. To find the saddle points of the Lagrangian, we can solve the following system of equations:

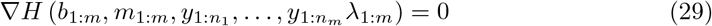

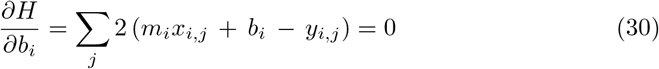

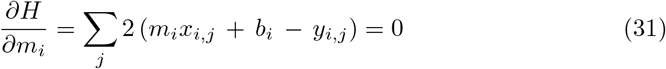

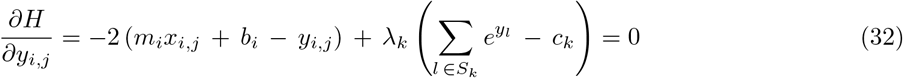

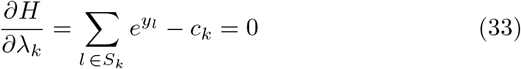

Solving this system of equation produces *m* and *b* values for each genome, and therefore can be used to determine abundances and PTRs for each genome. Like the gradient descent-based method described earlier in this paper, this can be modified to include *L*_1_ and *L*_2_ regularization, Poisson likelihoods, and amplicon bias. However, due to the numerical instability of this method, we decided to move away from using this approach.

### B Proposal for experimental validation

#### B.1 Minimum viable dataset

The minimum viable dataset to validate our method would need to consist of amplicon and WGS sequencing data verifiably derived from the same (not similar!) samples. Additionally, the sequences would need to meet the coPTR coverage requirements (5,000 WGS reads per OTU per sample). Based off of the accuracies reported in Fig. 2, we would also need more than 1,000 amplicon reads per OTU per sample.

To perform this comparison, we would repeat the procedure laid out for the DIABIMMUNE experiments in Sec. 3.5.2: produce estimates for coPTR and aPTR and compute the correlation between matching (OTU, sample) pairs of PTR estimates.

#### B.2 Resequencing existing samples

It is possible that existing samples, if properly frozen, may be sent to be sequenced using a different method without affecting the DNA content in the sample, yielding an appropriate comparison without advance planning.

Although any hypervariable region can be used for analysis, full-length 16S RNA sequencing allows for the greatest number of distinct OTUs to be analyzed. PacBio sequencing would be required to accommodate the length of the full 16S RNA sequence. Amplicon resequencing experiments would also benefit from greaterthan-average read coverage to ensure that the threshold of 1,000 reads per OTU per sample is met for lower-abundance species.

#### B.3 Ground-truth growth rates

It is preferable to work with a known, ground-truth growth rate. For this, in addition to the requirements laid out in Sec. B.1, we need ground-truth growth rates. This would best be produced by performing optical density measurements on microbial cultures at each sampling time. Combining optical densities with relative abundances (which can be inferred from amplicon data easily) yields absolute abundances; dividing these by the interval of time between samples yields an average growth rate.

Such datasets are sometimes generated for microbial time-series data, e.g. in the Lotka Volterra modeling literature, to transform relative into absolute abundances. To our knowledge, they have never been sequenced using WGS and amplicon methods in parallel.

In the presence of known ground truth growth rates, the same processing as in Secs. 2.5.3 and B.1 can be applied, but computing the correlations of each estimate with the true PTRs. Unlike these comparisons, these correlations must be computed per OTU, as the relationships between PTR and true growth rate vary from species to species.

#### B.4 Chemostat experiment

Rather than rely on observing growth rates, we can also follow the method of [8] and control microbial growth rates by use of a chemostat. The sequencing and processing requirements would be equivalent to the above sections.

#### B.5 Custom primers

One of the major impediments to computationally inferring PTRs in amplicon data is the large number of identical sequences (summarized in the **S** matrix). This results from the universal primer design, which only amplifies DNA inside of the 16S RNA gene, which is much more concerned than the surrounding genomic regions. Custom primers that extend past the start and end of the 16S gene would yield distinct sequences for each copy of the 16S RNA gene, obviating the need for the **S** matrix. For a clonal species, this would allow us to measure the correlation between genomic position and average coverage at different growth rates, the simplest possible validation of the idea behind our method.

## Notes

http://www.github.com/pchlenski/aptr

### Competing Interest Statement

The authors have declared no competing interest.

