## Supplementary material for "Signals of microbial growth learned from single amplicon samples": MLCB 2023 Extended Abstract

Philippe Chlenski[1*], Deirdre Ricaurte[1], Itsik Pe'er[1]
[1] Columbia University

**Introduction.** Whole-genome shotgun (WGS) read coverage along a bacterial chromosome decays log-linearly in both directions from the origin of replication (OOR) to the replication terminus (RT) [8, 5]. The ratio of coverages at these extremes, known as the "peak-to-trough ratio" (PTR), is correlated with microbial growth rates. With sufficient coverage, PTR estimates can be used to learn species growth rates in a metagenomic sample. However, many metagenomic studies eschew WGS reads in favor of amplicon sequencing of the 16S rRNA gene. While this approach reveals the taxonomic composition of a sample cost-effectively [2] and has a wealth of taxonomic reference data [4, 17], this data suffers from several complications [6, 1, 19, 10, 11, 12]. Additionally, it renders traditional PTR-based analysis impossible by concentrating all read coverage at 16S genes. In this work, we look for multicopy diverged (MCD) 16S sequences to perform a modified form of PTR analysis appropriate to 16S amplicon sequence data.

**Reference data collection.** We filter PATRIC reference genomes [3, 13] to those with MCD 16S genes:

1. Trim 16S sequences by the study primers
2. Remove genomes without a contig containing MCD 16S sequences
3. Remove genomes that share a 16S sequence with any genomes removed in step (2)
4. Repeat steps 2–3 until convergence
5. Keep only genomes with a single known OOR.

**Generative model.** Let $S, T, G$, and $N$ be the number of samples, taxa, genes, and distinct 16S rRNA gene nucleotide sequences in an experiment, indexed by $s, t, g$, and $n$. We consider the coverage pattern for a taxon $t$ in sample $s$: during exponential growth, the log-PTR for taxon $t$ in sample $s$ is $R_{t,s} = \log(\rho_{t,s}/\tau_{t,s})$, where $\rho_{t,s}$ is the replication time and $\tau_{t,s}$ is the generation time. If $A_{t,s}$ is the coverage at the OOR, the relative distance $\delta_t(l)$ from $l$ to the OOR $l_{0,t}$ on a chromosome of size $L_t$, and the expected coverage $C_{t,s}(l)$ are given by:

$$\delta_t(l) = \frac{\min\left(|l - l_{0,t}|,\ L_t - |l - l_{0,t}|\right)}{L_t} \tag{1}$$

$$\mathbb{E}\left[C_{t,s}(l)\right] = A_{t,s}\exp(-R_{t,s}2\delta_t(l)). \tag{2}$$

We aggregate these abundances and log-PTRs into $T \times S$ matrices $\mathbf{A}$ and $\mathbf{R}$. Letting $l_{g,t}$ be the locus at which gene $g$ starts in taxon $t$, we define an $N \times G$ sequence-sharing matrix $\mathbf{S}$, a $G \times T$ distance matrix $\mathbf{D}$, and a $G \times T$ membership matrix $\mathbf{M}$, and use them to express the $N \times S$ matrix $\hat{\mathbf{Y}}$ of expected read counts:

$$S_{ng} = \begin{cases} 1 & \text{Gene } g \text{ has sequence } n \\ 0 & \text{otherwise} \end{cases};\ D_{gt} = \begin{cases} \delta_t(l_{g,t}) & \text{Taxon } t \text{ has gene } g \\ 0 & \text{otherwise} \end{cases};\ \mathbf{M} = \operatorname{sign}(\mathbf{D}) \tag{3}$$

$$\hat{\mathbf{Y}}(\mathbf{A}, \mathbf{R}) = \mathbf{S}\left(\exp(-\mathbf{D}\mathbf{R}) \odot \mathbf{M}\mathbf{A}\right). \tag{4}$$

For unnormalized count data, it is natural to adopt a Poisson likelihood function. Since the entries of $\hat{\mathbf{Y}}$ already correspond to expected read counts, they can be used directly as the parameter of a Poisson likelihood function. Letting $\mathbf{Y}$ refer to the $N \times S$ matrix of empirical OTU read counts,

$$\ell(\mathbf{A}, \mathbf{R}) = \ln\left(\prod_{n=1}^{N}\prod_{s=1}^{S}\frac{\exp\left(-\hat{\mathbf{Y}}(\mathbf{A},\mathbf{R})_{ns}\right)}{\mathbf{Y}_{ns}!}\hat{\mathbf{Y}}(\mathbf{A},\mathbf{R})_{ns}^{\mathbf{Y}_{ns}}\right) = \frac{\left\|\hat{\mathbf{Y}}(\mathbf{A},\mathbf{R}) - \mathbf{Y}\odot\ln(\hat{\mathbf{Y}}(\mathbf{A},\mathbf{R})) + \ln(\mathbf{Y}!)\right\|_1}{NS}. \tag{5}$$

*

**Inference model.** Given a matrix of empirical observations $\mathbf{Y}$, matrices $\mathbf{S}$, $\mathbf{D}$, and $\mathbf{M}$ reflecting our knowledge of the distribution of 16S RNA sequences, and Equation 5, we can learn the PTRs and abundances of each organism. Specifically, we wish to find $\hat{\mathbf{A}}$, $\hat{\mathbf{R}}$ such that

$$(\hat{\mathbf{A}}, \hat{\mathbf{R}}) = \arg\min_{(\mathbf{A},\mathbf{R})} -\ell(\mathbf{A}, \mathbf{R}). \tag{6}$$

We solve this by gradient descent. Letting $\oslash$ represent element-wise division, we compute closed-form partial derivatives of $\ell$ as defined in Equation 5 with respect to $\mathbf{A}$ and $\mathbf{R}$:

$$\frac{\partial \ell(\mathbf{A}, \mathbf{R})}{\partial \mathbf{A}} = \mathbf{M}^T \mathbf{X}; \ \frac{\partial \ell(\mathbf{A}, \mathbf{R})}{\partial \mathbf{R}} = -\mathbf{D}^T \mathbf{X}; \ \mathbf{X} := \frac{\exp\left(\exp(-\mathbf{D}\mathbf{R}) \odot \mathbf{M}\mathbf{A}\right) \odot \mathbf{S}^T (1 - \mathbf{Y} \oslash \hat{\mathbf{Y}}(\mathbf{A}, \mathbf{R})}{NS} \tag{7}$$

**Further modifications.** The above model is sufficient to learn $\mathbf{A}$ and $\mathbf{R}$ for idealized data. In practice, it can be helpful to add $L_1$ regularization to $\mathbf{A}$ to encourage sparsity, to add $L_2$ regularization to $\mathbf{R}$ to encourage smaller PTRs, and to include an $N$-dimensional, learnable amplicon bias term $\mathbf{h}$. Letting $\alpha_1$ and $\alpha_2$ be new scalar hyperparameters controlling the regularization strength,

$$\hat{\mathbf{Y}}'(\mathbf{A}, \mathbf{R}) = \mathbf{h} \odot \hat{\mathbf{Y}}(\mathbf{A}, \mathbf{R}) \tag{8}$$

$$\ell'(\mathbf{A}, \mathbf{R}) = \ell(\mathbf{A}, \mathbf{R}) + \alpha_1 \|\mathbf{A}\|_1 + \alpha_2 \|\mathbf{R}\|_2. \tag{9}$$

**Bioinformatics pipeline.** We use VSEARCH[15] on raw WGS reads at an identity threshold of 1.00 (perfect sequence identity) to produce the empirical coverage matrix $\mathbf{Y}$. We then filter the sequences represented in $\mathbf{Y}$, keeping only those contained in the reference database and those corresponding to a genome with each MCD 16S. $\hat{\mathbf{A}}$ and $\hat{\mathbf{R}}$ are estimated according to Equations 6 and 7 using PyTorch [14].

**Results: synthetic data.** In synthetic datasets drawn by computing $\mathbf{Y} = \hat{\mathbf{Y}}$ using Equation 4 for known but held-out matrices $\mathbf{A}$ and $\mathbf{R}$, we found our method to have good performance retrieving PTRs starting at $10^5$ total reads in MCD sequences. We found no decrease in performance from increasing $T$, the total number of taxa considered in a single sample, or from switching from a negative Poisson log-likelihood to a mean squared error (MSE) objective. Including an amplicon bias term substantially improved performance in the presence of simulated amplicon bias and did not adversely affect performance when amplicon bias was not simulated.

**Results: comparison to coPTR.** We further compared our method to coPTR[5], the current state of the art in WGS-based PTR inference. First, we created synthetic datasets by drawing WGS reads at each locus using Equation 1, with reads in 16S regions tallied in $\mathbf{Y}$. We found that, so long as the $10^5$ read threshold identified in the previous section was met, our PTR and abundance estimates were comparable with coPTR. However, coPTR always had substantially more reads to work with and therefore outperformed our method. We then compared performance on the DIABIMMUNE dataset [22, 9, 20], where samples were sequenced using WGS and 16S methods, with relatively high 16S read coverage and number of samples. On this dataset, we found no correlation (Spearman $R = 0.01$) with coPTR estimates of PTR.

**Conclusions.** We present a novel extension of methods for PTR inference in WGS data to microbial relative-abundance data. We leverage rich reference data to pinpoint genomic positions for 16S sequences and produce a complete model of sequence coverage. We use this model to learn PTRs and abundances for each species in the system. We provide a mathematical model with a PyTorch implementation and an accompanying bioinformatics pipeline to preprocess metagenomic reads.

We demonstrate the effectiveness of our model on simulated data and test its robustness to number of genomes, total read coverage, amplicon bias, and loss function specification. We also report on an attempt to test our model on real data which found no correlation with WGS-based PTR inference methods. Future work should focus on identifying high-quality datasets for further comparison, validating this model with *in vitro* systems with known growth rates, and exploring confounding by evolutionary strategy [7, 18, 21, 16].
